# The knowledge and practice towards COVID-19 pandemic prevention among residents of Ethiopia: An Online Cross-Sectional Study

**DOI:** 10.1101/2020.06.01.127381

**Authors:** Daniel Bekele, Tadesse Tolossa, Reta Tsegaye, Wondesen Teshome

## Abstract

**Background:** The Novel coronavirus disease-2019 (COVID-19) is now the international concerns and a pandemic, since the World Health Organization declared as the outbreaks. The objective of this study to assess the prevention knowledge and practices towards the COVID-19 among the residents of Ethiopia.

**Methods:** An online cross-sectional study was conducted among the sample of Ethiopian residents via social platforms of the author’s network with the popular media such as Facebook, in Ethiopia from the April 15-22, 2020 and successfully recruited 341 responses. The snowball sampling was employed to recruit the participants. The data were analyzed using STATA version 14. Descriptive statistics were used to summarize the level of knowledge and practices.

**Results:** The majority of the respondents (80.35%) were male. The overall prevention knowledge of the participants towards the novel coronavirus (COVID-19) was high. About 91.2% of the participant was heard about the novel coronavirus disease and Social Medias’ were the main source of the information. About 90% of the participants had a good prevention knowledge of maintaining social distance and frequent hand washing. The practices of the participants towards the COVID-19 prevention were very low. Out of 341 participants, only 61% and 84% of the participants were practicing social distance and frequent hand washing, respectively.

**Conclusions:** Majority of the participants had knew the ways of protecting themselves from the novel coronavirus. But, there was deficiencies of changing these prevention knowledge to practice. This indicates that there was gap between knowledge and implementation of prevention strategies in the community. The concerned body should focus on providing education for the community regarding the implementation of the prevention knowledge to practice.

## Introduction

Coronavirus is one of the major pathogens that mainly target the respiratory system of humans. Previous outbreaks of coronaviruses were recorded in history as severe acute respiratory syndrome (SARS)-CoV and the Middle East respiratory syndrome (MERS)-CoV (GeoPoll 2020). The new coronavirus identified as the cause of the acute respiratory disease since the end of December 2019, later labelled as SARS-CoV2 by World Health Organization is a different strain of coronavirus from SARS and MERS coronaviruses. The difference is genetic make-up, clinical presentations, case fatality and the rate of spread across the world. SARS-CoV2, the virus that causes coronavirus disease 2019 (COVID-19), become the newest virus to cause global health fear (WHO, 2020; Nuwagira and Muzoora 2020).

The COVID-19 symptoms can range from mild (or no symptoms) to severe illness and are mainly characterized by fever, dry cough, dyspnea, headache, sore throat and rhinorrhea and sometimes hemoptysis (Adhikari et al. 2020, Cascella, Rajnik et al. 2020). The main route of transmissions are close contact (about 6 feet or two arm lengths) with a person who has COVID-19, respiratory droplets when an infected person coughs, sneezes, or talks and touching a surface or object that has the virus on it, and then by touching mouth, nose, or eyes (Guo et al. 2020).

Currently, more than 2.2 million people were getting infected, and more than 152 thousands died of coronavirus globally as of 19 April 2020 (Roy et al. 2020), and the figures are still increasing rapidly. In Africa, almost all countries have now confirmed cases and the number of deaths is increasing. If the spread of the disease is not well managed, its impact on the African economy will be extensive. Africa’s already fragile health systems, coupled with a high burden of Covid-19 would cost the continent greatly. The speed with which countries can detect report and respond to outbreaks can be a reflection of their wider institutional capacity (Ikhaq, Riaz et al. 2020, Olapegba et al. 2020). The threat of COVID-19 to health systems in Africa can be compared with a metaphor which says “the eye of the crocodile”; which means in the lake, only the eyes of the crocodile is visible on the surface while the rest of the body is immersed in water. In this viewpoint, the eyes represent the public health preparedness while the body of the crocodile represents Africa’s fragile health systems (Paintsil, 2020).

According to the Ministry of Health, Ethiopia, the number of infected cases reached 117 and 3 deaths as of 24 April 2020. In a developing country like Ethiopia, where trained human resources and equipment for the treatment of COVID-19 are scarce, working on prevention of the viral spread should be a priority and feasible intervention. Government of Ethiopia has declared a state of emergency to stop the spread of the disease. These include implementation of staying home as much as possible, avoiding close contact with others, cleaning and disinfecting frequently touched surfaces, washing hands often with soap and water for at least 20 seconds, or using hand sanitizer containing at least 60% alcohol (Cascella et al. 2020). For successful implementation of these preventive measures, knowledge of the community about coronavirus disease (Covid-19) and practice of its preventive measures have to be optimum. However, the level of knowledge and practice of the people is not well understood. Thus, this study is intended to identify the level of knowledge and practice towards Covid-19 prevention among residents in Ethiopia.

### Methodology

This study employed an online cross-sectional survey by using the Google Forms to collects the data from the respondents regarding their prevention knowledge and practice towards the COVID-19 pandemic from April 15-22, 2020. Since, the first cases of the COVID-19 confirmed in Ethiopia on March 13, 2020 the government declared the state lockdown on April 5, 2020. These made difficult to conduct the community or institutional based survey. Depending on this to select the respondents from the population snowball sampling technique was employed through the author’s network with residents on the popular social media such as; Facebook, Telegram, and Email in Ethiopia. The link of the questionnaire was posted by the author’s on the above mentioned Social Media and to fulfill the Terms and Conditions of website regarding the responses of respondents the following questions was added at the end of the questionnaire on the Google Forms, said that “By submitting this form, do you agree to the Terms and Conditions of Google Forms” and if they agreed with this Terms and Conditions of the Google Forms their responses can be submitted. So, we are confidential that our data collection method was compiled with the Terms and Conditions of the Google Forms website. The questionnaire contained both an open ended and closed ended questions that focused on the respondents Socio-demographic, prevention knowledge, and practice towards the COVID-19. The inclusion criteria of the participants used in this study was: be able to read and understand English language, have Ethiopian nationality, and be aged ≥ 18 years. Prior proceeding to fulfill the questionnaire objective of the study, the informed consent of participation, the declaration of anonymity and confidentiality of their response was briefed in well manner in the questionnaire (Roy et al. 2020).

### Measurements and Data Management

The prevention knowledge and practice towards the COVID-19 were measured based on the WHO (2020) Survey Tool and Guidance (WHO, 2020b). This study was employed descriptive statistics to summarize the prevention knowledge and the practice of their respondents towards the Novel coronavirus pandemic(WHO, 2020a). The questions about prevention knowledge had 7 items as presented in Table **2**, and the questions about the respondent’s practice of prevention knowledge had 6 items as presented in Table **3** and the rest of the four questions was about the respondent’s Socio-demographic information. All the questions contained the category of (“Yes”, “No”, “Don’t know”). The prevention knowledge and the practice level was assessed by assigning one point for each correct answer and the prevention knowledge indicated by two category: poor for (< 5 of 7 items) and good for (≥ 5 of 7 items). Specially regarding of the practice of prevention knowledge the respondents was asked about going crowded place, wearing mask in public, maintain the social distance, hand washing, avoid handshaking, and obeying the government restriction. The practices of participants was indicated by two category: poor practice for (< 4 of 6 items) and good practice for (≥ 4 of 6 items).

The collected data was checked manually for its completeness. The data was coded and edited in Epi-Info version 3.5.1 and exported to STATA version 14.0 for further statistical data analysis. The descriptive statistics was employed to summarize the data of this study.

### Ethical Approval and Considerations

The study was conducted by preparing the online survey and the investigators informed the study participants about the general objective of the study and that their answer were remain anonymous to ensure the confidentiality of the information they provided. So, their consent of participation was informed prior to proceeding to fill questionnaire to every respondents. The questionnaire was designed to be anonymous and the result did not identify the personality of the respondents rather presented in the aggregated statistics.

## Results

### Socio-demographic characteristics of the respondents

This study was involved among 341 respondents in which the majority (80.35%) were males. More than the two thirds of the respondents were in the age group of 18-32 and had a level of education of college and above. Of all respondents, more than half of them were single in marital status, and about the two thirds of them were urban residents while 16.3% were rural residents (Table 1).

**Table 1.**
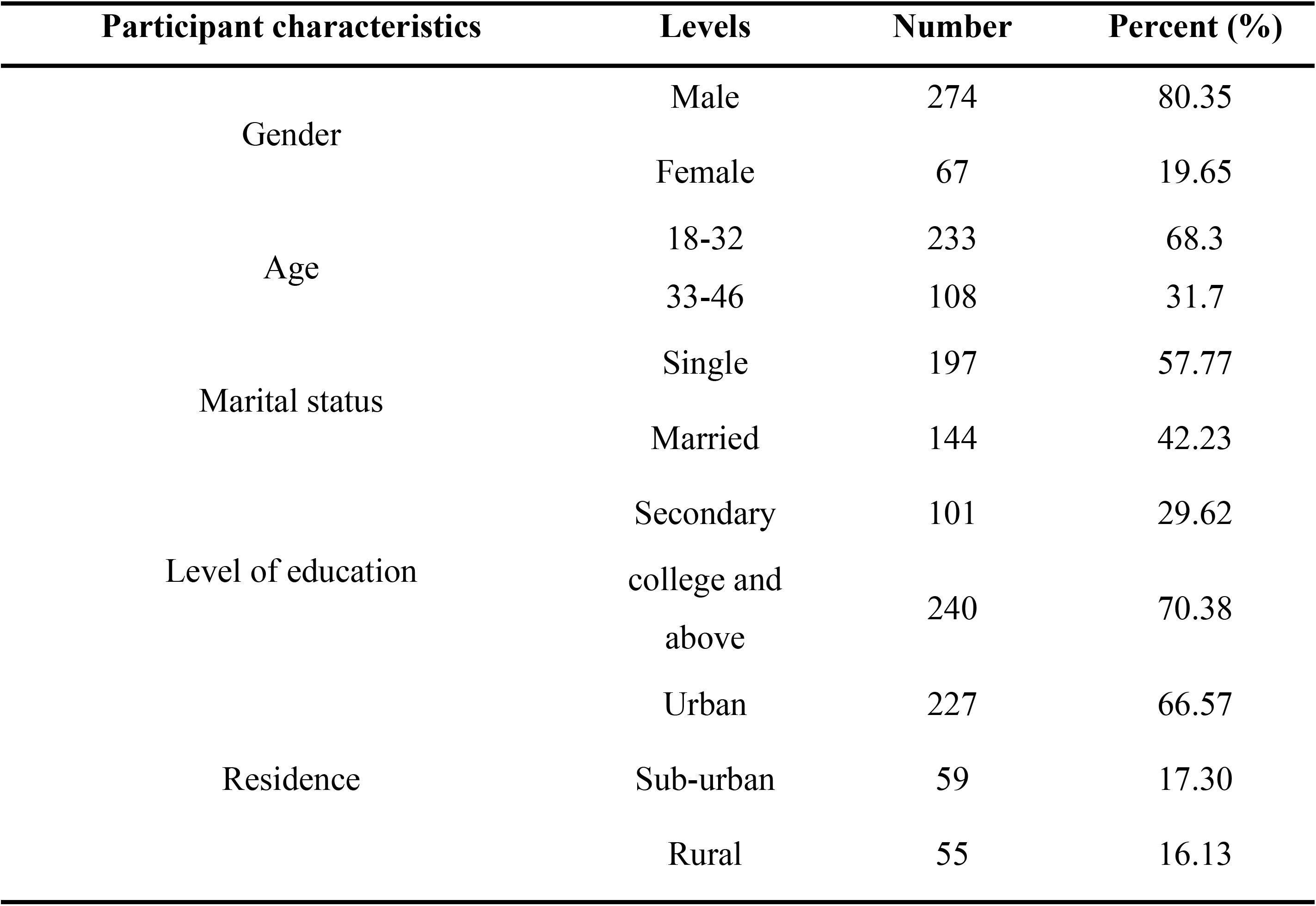
Socio-demographic characteristics of respondent’s to assess the prevention knowledge, practice of among community towards COVID-19 pandemic in Ethiopia (N=341).

### The Prevention Knowledge of the respondents towards the COVID-19

In the current study, from all respondents, the majority (91.8%) had heard of Covid-19. Though the majority of the respondents thought to avoid touching nose, eye, and face with unwashed hand protects from getting of the COVID-19, and about 11.4% of the respondents do not agree with this preventive measure. The 286 (83.9%) out of all the respondents thought that wearing mask can protects from getting of the COVID-19 and rest of 16.1% of the respondent’s do not knew wearing a face mask can protect from getting of the COVID-19. Majority, three-fourth of the respondents reported that avoiding hugging with people can protect from getting the infection of COVID-19 and 26.7% out of all respondents thought that drinking a lot of water can protect from getting infection of the COVID-19. More than 93% of the respondents reported that maintaining social distance can protect from getting of the COVID-19 and 90% of them thought that frequent hand washing for 20 seconds can protect from getting infection of the COVID-19 (Table 2)

**Table 2:** Frequency distribution of respondents’ knowledge on prevention Covid-19, Ethiopia (N =341).

| Knowledge questions on prevention of Covid-19 | Yes | No |
| --- | --- | --- |
|  | N (%) | N (%) |
| Have you heard of COVID-19? | 313(91.8) | 28(8.2) |
| Do you think avoid touching nose, eye, and face with unwashed hand protect COVID-19? | 302(88.6) | 39(11.4) |
| Do you think wearing mask protect from COVID-19? | 286(83.9) | 55(16.1) |
| Do you think avoid hugging with people protect from COVID-19? | 263(77.1) | 78(22.9) |
| Do you think drinking a lot of water can protect from COVID-19? | 91(26.7) | 250(73.3) |
| Do you think maintaining social distance can protect from COVID-19? | 320(93.8) | 21(6.2) |
| Do you think frequently washing hand for 20 second can protect COVID-19? | 307(90.0) | 34(10.0) |

In this study, the urban area residents have a good knowledge of the COVID-19 prevention through avoiding hugging with peoples than the respondents live in suburban and rural area, respectively (Figure 1).

**Figure 1:**
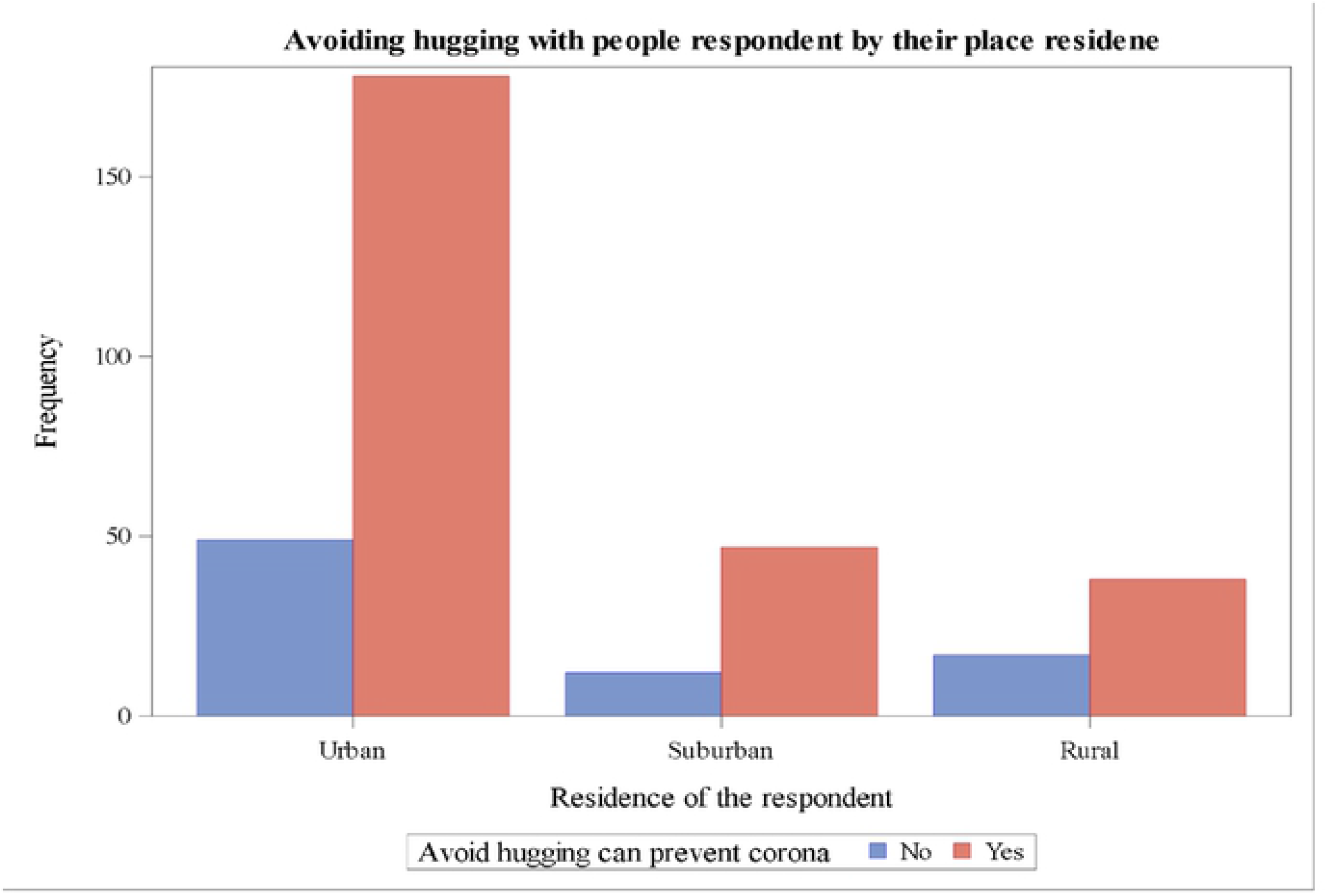
The proportion of respondents’ knowledge that avoiding hugging with people can protect from ting of COVID-19 pandemic by their place of residence.

The majority of the respondents who had highest educational level knew that avoiding touching of eye, face, and nose with unwashed hand can reduce the risk of getting infected with the Novel Coronavirus (COVID-19) (Figure 2).

**Figure 2:**
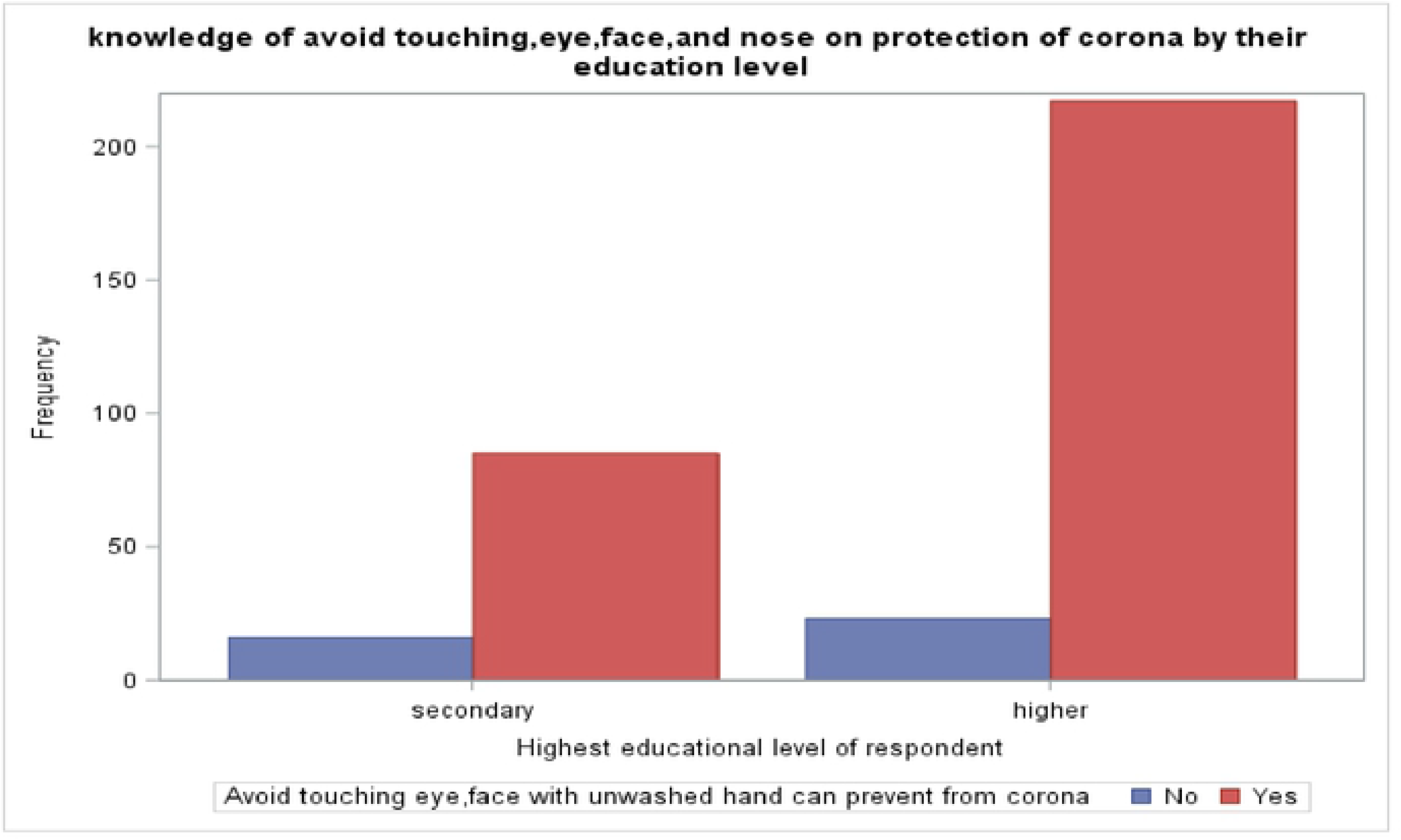
The proportion of respondent knowledge on avoid touching eye, nose, and face can protect from Novel Coronavirus (COVID-19) by their level of education.

The majority of the respondents who had highest educational level reported that they knew that maintaining social distance can reduces the risk of getting with the Novel Coronavirus (COVID-19). On other hand, respondents with secondary level of education and below had less knowledge on maintaining social distance can reduce the risk of getting the COVID-19 (Figure 3).

**Figure 3:**
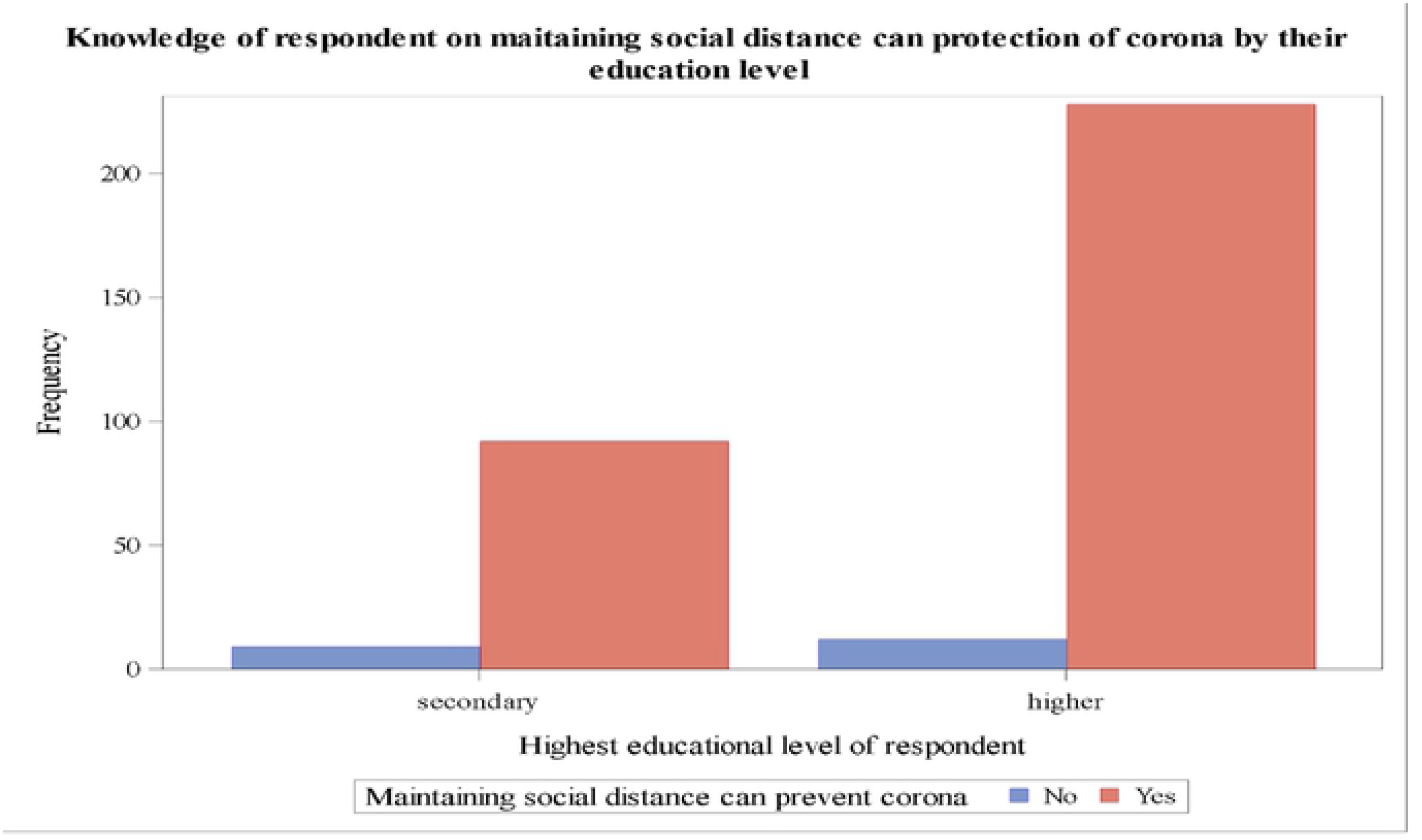
The proportion of respondent knowledge on maintaining social distance can protect from Novel Coronavirus (COVID-19) by their level of education.

### The respondents practices on the prevention of the COVID-19

In this study, about 77.4% of the respondents were not obeying government restrictions, whereas; only about 22.6% of the respondents were obeying the government restriction on the COVID-19 prevention. About 137(40.2%) of the respondents were a still going to crowded places and about 76% of them were not wearing a face masks when leave the home. About 61.1% of the respondents were keeping themselves two meters from another person’s in the public place while 39.9% of the respondents were not experiencing social distance in the public place. Majority (84.5%) of the respondents wash their hand when they go to the public place and 20.8% of the respondents are still making the handshake with people when they go to the public place (Table 3).

**Table 3:** Frequency distribution of respondents response on their practice towards Novel Coronavirus (COVID-19) pandemic in Ethiopia (N =341)

| **Practice related questions towards Covid-19 prevention** | **Yes**<br>**N (%)** | **No**<br>**N (%)** |
| --- | --- | --- |
| Are you obeying government restriction on COVID-19 prevention strategies? | 264(77.4) | 77(22.6) |
| Are you going to public or crowded place after the COVID-19 pandemic confirmed? | 137(40.2) | 204(59.8) |
| Did you wear mask when you leave your home? | 82(24.0) | 259(76.0) |
| Did you keep yourself 2 meter from another person when you go to public? | 210(61.6) | 131(38.4) |
| Did you make a handshake with a person when you go to the public? | 71(20.8) | 270(79.2) |
| Did you wash your hand at the place you went to or in the public? | 288(84.5) | 53(15.5) |

Majority of the respondents who live in the urban area had a good implementation of prevention knowledge towards the COVID-19 pandemic through wearing a face mask when they went to the public place as compared to the respondents live in suburban and rural area, respectively (Figure 4).

**Figure 4:**
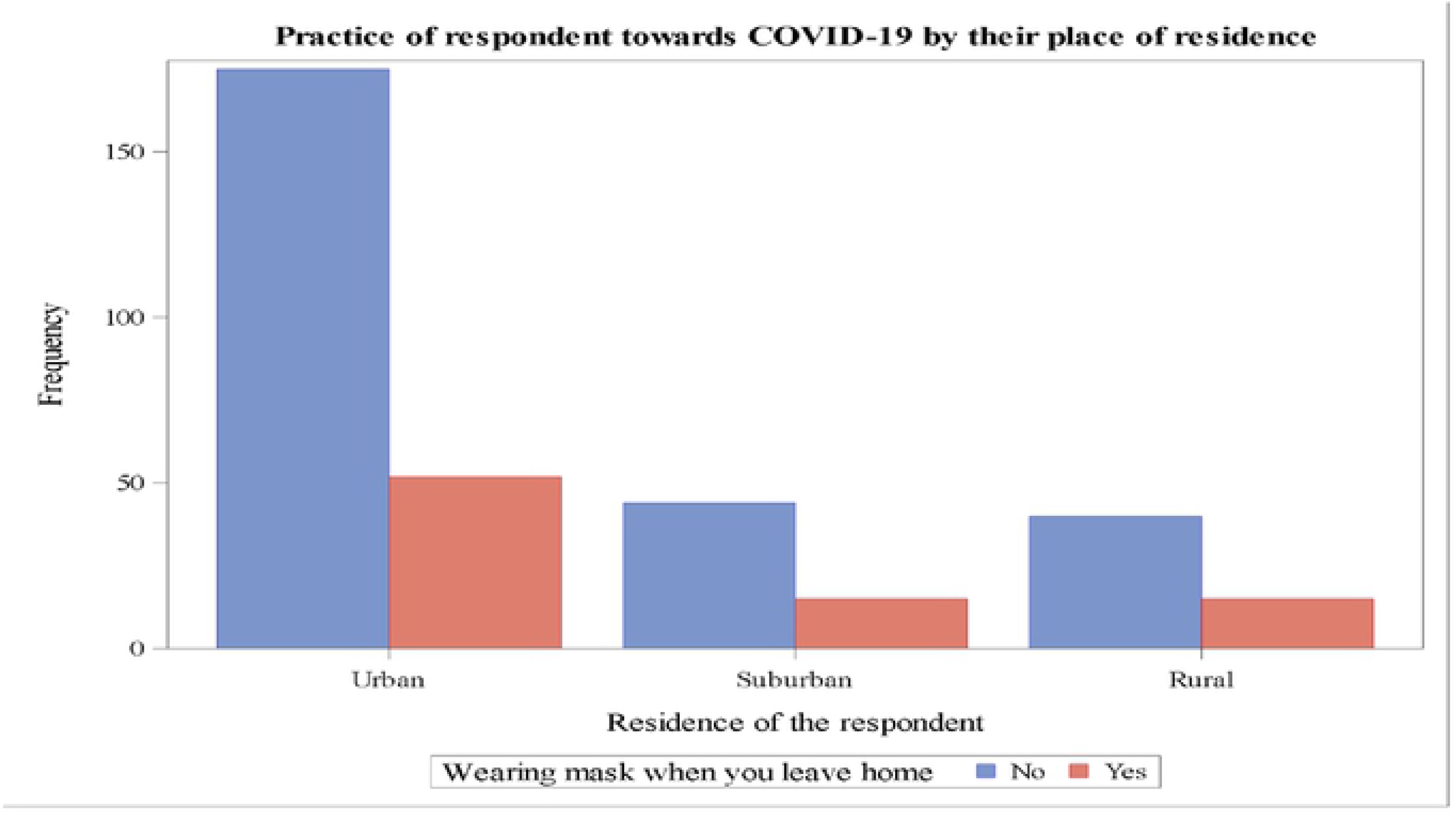
The practice of respondent towards the Novel Coronavirus (COVID-19) by their place of residence.

Those respondents with highest educational level had a good practices of prevention knowledge towards the COVID-19 pandemic by maintaining the social distance (i.e., at least keeping yourself two meters from another person) as compared to the respondents with secondary educational level (Figure 5).

**Figure 5:**
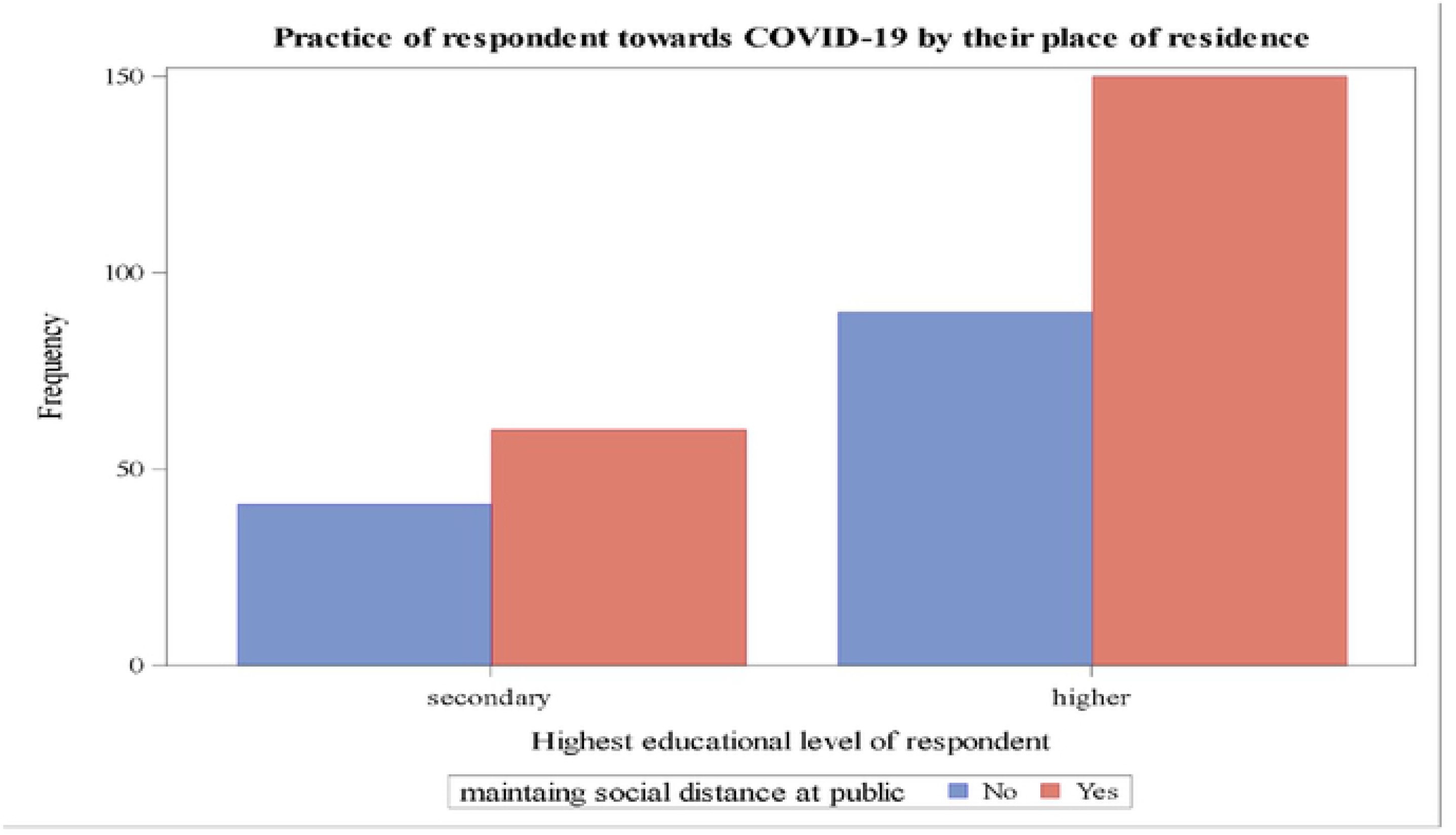
The practice of respondent towards the Novel Coronavirus (COVID-19) by their level of education.

The mean score and standard deviation of the respondents prevention knowledge of the COVID-19 pandemic was 5.52 ±1.11 and the mean score and standard deviation of the respondents on the practice of prevention knowledge of the COVID-19 pandemic was 3.09 ± 1.06 (Table 4).

**Table 4:** The prevention knowledge and practice score of the respondents towards the COVID-19 pandemic (N=341).

| Variables | Mean ± SD | Score range |
| --- | --- | --- |
| Knowledge score on prevention Covid-19 | $5.52 \pm 1.11$ | Range (0-7) |
| Practice score on prevention of Covid-19 | $3.09 \pm 1.06$ | Range (0-6) |

In this study, the knowledge score of males was greater than the overall mean score while the knowledge score of females was below the mean score. Respondents aged 18 to 32 had knowledge score above the mean but lower than mean score in the practice of Covid-19 prevention. Respondents with a single in marital status, education level of college and above and urban residents had a better score of knowledge and practice, above the mean score, as compared to their counterparts (Table 5).

**Table 4.** Demographic characteristics of respondents by their knowledge score and practice score towards Novel Coronavirus (COVID-19) prevention in Ethiopia (N=341)

| Characteristics | | Number | Percent | Knowledge score<br>(mean ±SD) | Practice score<br>(mean ±SD) |
| --- | --- | --- | --- | --- | --- |
| Gender | Male | 274 | 80.35 | 5.63(1.02) | 3.12(1.01) |
|  | Female | 67 | 19.65 | 5.05(1.34) | 2.96(1.25) |
| Age | 18-32 | 233 | 68.3 | 5.55(1.09) | 3.07(1.08) |
|  | 33-46 | 108 | 31.7 | 5.45(1.15) | 3.13(1.01) |
| Marital status | Single | 197 | 57.77 | 5.59(1.08) | 3.20(1.02) |
|  | Married | 144 | 42.23 | 5.41(1.14) | 2.94(1.09) |
| Level of education | Secondary education | 101 | 29.62 | 5.28(1.29) | 3.03(1.16) |
|  | College and above | 240 | 70.38 | 5.62(1.01) | 3.12(1.02) |
| Residence | Urban | 227 | 66.57 | 5.60(1.06) | 3.13(0.98) |
|  | Sub-urban | 59 | 17.30 | 5.6(1.14) | 3.07(1.15) |
|  | Rural | 55 | 16.13 | 5.05(1.17) | 2.98(1.24) |

As we have seen from below (Figure 6) the most main source of information the participant ever heard about the COVID-19 pandemic were social media and television.

**Figure 6:**
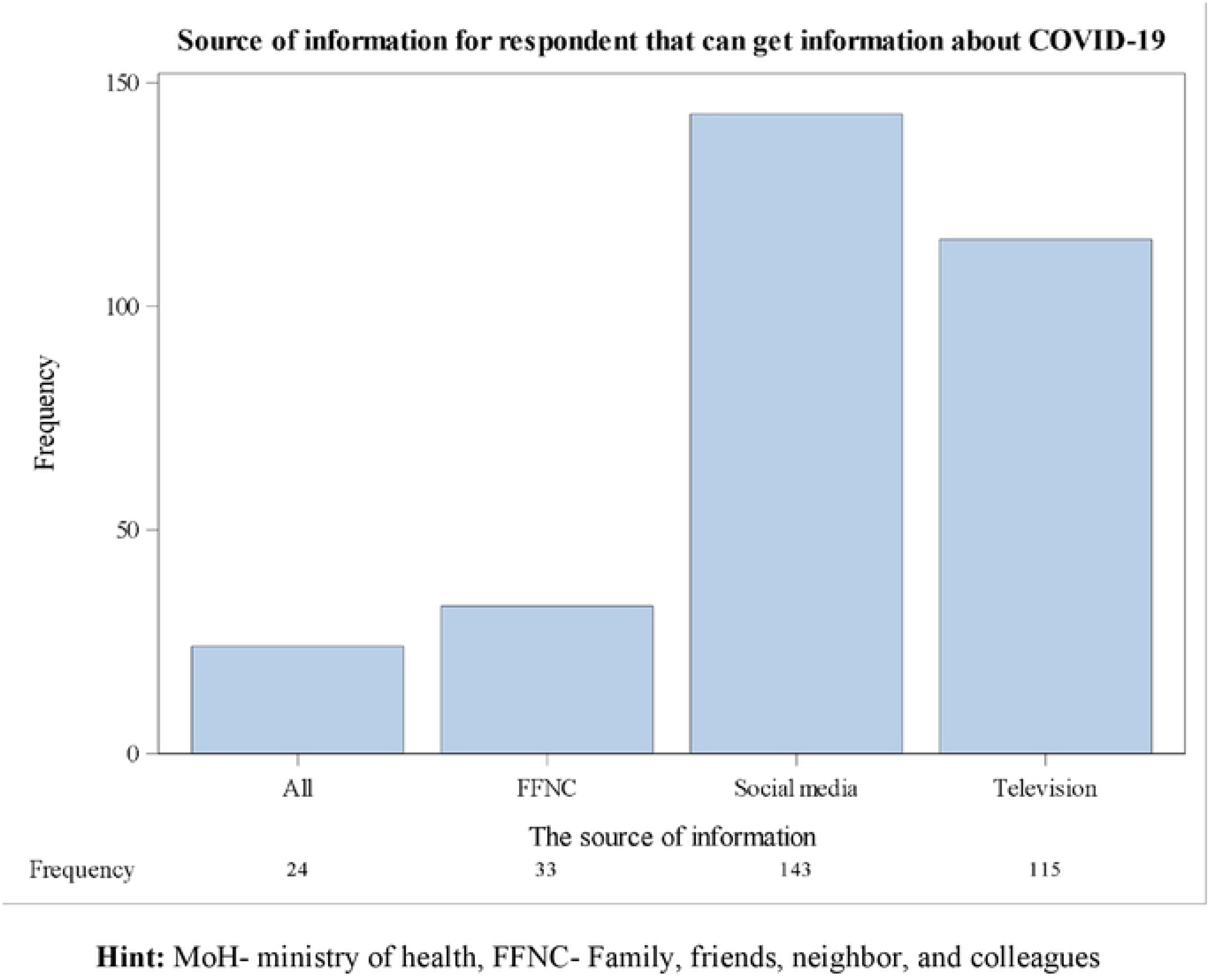
The source of information where respondent hear about Novel Corona virus (COVID-19) pandemic in Ethiopia (N=341)

## Discussion

This study was the first study to assess the prevention knowledge and practices of prevention knowledge towards the COVID-19 prevention among the residents of Ethiopia. Accordingly, the prevention knowledge of the respondents towards the novel coronavirus was high. Because of the government of Ethiopia was taking a lot of measurements after the first cases of the COVID-19 was confirmed and the measurement include the social media campaigns, awareness creation about the diseases, the closing school and university, the closing religious places, and declaring the state lockdown was used to minimize the transmission root of the diseases.

Based on knowledge score of the participants, overall prevention knowledge towards the COVID 19 pandemic was 78.8%. It is lower than the study conducted in China among health care workers in which the knowledge of HCW towards COVID 19 prevention was high (89%) (Zhou et al. 2020). Furthermore, this finding is lower than the finding conducted in Pakistan in which the knowledge towards COVID 19 prevention was 92.3% (Saqlain et al. 2020). This discrepancy could be due to the difference in the study population. The previous studies were conducted among health professional to assess their knowledge level, but the current study is conducted among the general population. This is due to the fact that health professional has more exposure to the information regarding the disease.

The finding of our study was far higher than the study conducted in Nigeria among pregnant women. The result showed that the knowledge of pregnant women regarding to the preventive mechanism of COVID 19 was 60.9% (Nwafor et al. 2020). The difference might be due to the time of the study and the population participated in the study. In a study conducted in Nigeria, the disease was declared as a pandemic for the first time while our study was conducted after the population provided with awareness about the disease. In addition, in the previous study, pregnant women were participated, while in the current study residents were participated. Despite the knowledge level of Ethiopia population towards coronavirus prevention was high, the practice toward the disease prevention was poor. This could be due to various factors including the availability of personal protective in Ethiopia is inadequate, the economic status of the population is substandard and their life is depending on daily activity.

In our study, more than 90% of the population had a good knowledge with regards to maintaining social distance and frequently washing hand for greater than 20 seconds. These are the two main strategies which were recommended by WHO to prevent the transmission of the disease (WHO, 2020c). Even the knowledge regarding social distancing and frequent hand washing was good, the practice of experiencing social support and frequent hand washing is low when compared to their knowledge level. In our study, only 61% and 84% of participants practice social distancing and hand washing, respectively. Furthermore, around 88% of participants have good knowledge that wearing a mask can protect us from COVID 19. Similarly, 82 % of participants were practiced to worn mask to prevent COVID 19 in Ethiopia.

In our finding, around 91.2% of Ethiopian Population had heard about Novel coronavirus at the time of the study. It is comparable with a study conducted in Pakistan in which more than 90% of medical students had heard about the disease (Ikhaq et al. 2020). Furthermore, this finding is in line with a study conducted in Nepal in which 91.6 % of population aware all the clinical features of Coronavirus (Vaidya et al., 2020). A survey conducted in three countries of Africa (Nigeria, South Africa and Kenya) showed that greater than 94% of the population were heard the coronavirus (GeoPoll, 2020).

The main source information for the participants was social media, followed by the Television, which is in line with the multinational study done in India (Kamate, et al. 2020). But it is in contradict with a study done in Nigeria in which the main source of information was mass media (Olapegba et al., 2020). It is obvious that during the development of a newly emerging disease like coronavirus, there might not be enough published literature on local or international journals. In such cases, social medias’ such as Facebook and telegram plays a major role in providing up-to-date information for the population. In addition, television and radio are the main source information next to social media, but it depends on various factors like availability of light, cost, and mostly available in urban town. In Ethiopia, most of the population live in rural/remote area, and the availability of Television are very low because the coverage of the electric light is low, hence, the population are forced to use easily accessible material such as mobile data.

### Strength and limitation of the study

The strength of this study was it is the first study to assess the prevention knowledge and practice of prevention knowledge towards the Novel coronavirus diseases among the residents of Ethiopia. There are some limitations to this research. First, questions related to knowledge and practices are not validated. The second and the major limitation of this study was the data collected by an online distribution of the survey through the Social Media and only individual with the access of Social Media and internet can participate in the study and this study cannot represent the overall population of the country.

### Conclusions

In this study, almost the prevention knowledge of the respondent towards the COVID-19 pandemic was high. But, the practice of the respondents prevention knowledge to the implementation very low and the practices score of the respondent was (mean=3.09), which very low and which indicates that the practices the respondents towards the COVID-19 pandemic was poor. As far as the result of this study shows the respondents had a good prevention knowledge of the COVID-19 pandemic. To tackle the spreads of the COVID-19 pandemic in the community the prevention knowledge of the community should be implemented into the practices and the stakeholders should play their significant contribution by teaching the community in ways to implementing the prevention knowledge to combat the spread of the Novel coronavirus pandemics.

## Abbreviations

FFNC: Family, friends, neighbor, and colleagues
MoH: Ministry of Health
SARS: Severe Acute Respiratory Syndrome
MERS: Middle East Respiratory Syndrome
WHO: World Health Organization

## Declaration

### Consent for publication

Not applicable

### Availability of data and materials

All the relevant datasets used in this study was available in the manuscript and in its supporting information files.

### Authors’ Contribution

DB involved in the developing initial idea of the research, designing of the questionnaires on web, and statistical analysis of the result. TT, RT and WT involved in writing the body of the research. All authors involved in developing the initial drafts of the manuscript, revising subsequent drafts and prepared the final draft of the manuscript. All authors read and approved the final draft of the manuscript

## Reference

Adhikari SP., Meng S, Wu YJ, Mao YP, Ye RX, Wang QZ, Sun C, Sylvia S, Rozelle S and Raat H (2020). Epidemiology, causes, clinical manifestation and diagnosis, prevention and control of coronavirus disease (COVID-19) during the early outbreak period: a scoping review. Infectious diseases of poverty, 9(1): 1–12.

Cascella M, Rajnik M, Cuomo A, Dulebohn SC and Napoli RD (2020). Features, evaluation and treatment coronavirus (COVID-19). Statpearls [internet], StatPearls Publishing.

GeoPoll (2020). A study of the knowledge and perceptions of coronavirus (COVID-19) in South Africa, Kenya, and Nigeria.

Guo YR, Cao QD, Hong ZS, Tan YY, Chen SD, Jin HJ, Tan KS, Wang DY and Yan Y (2020). The origin, transmission and clinical therapies on coronavirus disease 2019 (COVID-19) outbreak-an update on the status. Military Medical Research 7(1): 1–10.

Ikhaq A, Riaz HBE, Bashir I and Ijaz F (2020). Awareness and Attitude of Undergraduate Medical Students towards 2019-novel Corona virus. Pakistan Journal of Medical Sciences 36 (COVID19-S4).

Kamate SK, Sharma S, Thakar S, Srivastava D, Sengupta K, Hadi AJ, Chaudhary A, Joshi R and Dhanker K (2020). Assessing Knowledge, Attitudes and Practices of dental practitioners regarding the COVID-19 pandemic: A multinational study. Dental and Medical Problems 57(1): 11–17.

Nuwagira E and Muzoora C (2020). Is Sub-Saharan Africa prepared for COVID-19? Tropical Medicine and Health 48(1): 1–3.

Nwafor J I, Aniukwu JK, Anozie BO and Ikeotuonye AC (2020). Knowledge and practice of preventive measures against COVID-19 infection among pregnant women in a low-resource African setting. medRxiv.

Olapegba PO, Ayandele O, Kolawole SO, Oguntayo R, Gandi JC, Dangiwa AL, Ottu IFA and Iorfa SK (2020). A Preliminary Assessment of Novel Coronavirus (COVID-19) Knowledge and Perceptions in Nigeria. medRxiv.

Paintsil E. (2020). COVID-19 threatens health systems in sub-Saharan Africa: the eye of the crocodile “The journal of clinical investigation

Roy, D., S. Tripathy, S. K. Kar, N. Sharma, S. K. Verma and V. Kaushal (2020). Study of knowledge, attitude, anxiety & perceived mental healthcare need in Indian population during COVID-19 pandemic. Asian Journal of Psychiatry: 102083.

Saqlain, M., Munir MM, Rehman SU, Gulzar A, Naz S, Ahmed Z, Tahir AH and Mashhood M (2020). Knowledge, attitude, practice and perceived barriers among healthcare professionals regarding COVID-19: A Cross-sectional survey from Pakistan. medRxiv.

Vaidya B, Bhochhibhoya M, Baral R and Nakarmi S (2020). Knowledge, Attitude and Practice towards COVID-19 among Patients with Musculoskeletal and Rheumatic Diseases in Nepal: A Web-based Cross-Sectional Study.

WHO (2020a). Coronavirus Disease 2019 (COVID-19) Situation Report - 60.

WHO (2020b) Survey Tool and Guidance: Rapid, Simple, Flexible Behavioural Insights on COVID-19. DK-2100 Copenhagen Ø, Denmark.

WHO (2020c). Critical preparedness, readiness and response actions for COVID-19: interim guidance, 22 March 2020, World Health Organization.

Zhou M, Tang F, Wang Y, Nie H, Zhang L, You G and Zhang M (2020). Knowledge, attitude and practice regarding COVID-19 among health care workers in Henan, China. Journal of Hospital Infection.

